# The effect of double biofeedback on functional hemispheric asymmetry and activity: a pilot study

**DOI:** 10.1101/2021.03.30.437721

**Authors:** Valeriia Demareva, Elena Mukhina, Tatiana Bobro

**Author notes:** **Publisher’s Note**: MDPI stays neutral with regard to jurisdictional claims in published maps and institutional affiliations.

## Abstract

In the current pilot study, we attempt to find out how double neurofeedback influences functional hemispheric asymmetry and activity. We examined 30 healthy participants (8 males; 22 females, mean age = 29; SD= 8). To measure functional hemispheric asymmetry and activity, we used computer laterometry in the ‘two-source’ lead-lag dichotic paradigm. Double biofeedback included 8 minutes of EEG oscillation recording with five minutes of basic mode. During the basic mode, the current amplitude of the EEG oscillator gets transformed into feedback sounds while the current amplitude of alpha EEG oscillator is used to modulate the intensity of light signals. Double neurofeedback did not directly influence the asymmetry itself but accelerated individual sound perception characteristics during dichotic listening in the preceding effect paradigm. Further research is needed to investigate the effect of double neurofeedback training on functional brain activity and asymmetry taking into account participants’ age, gender, and motivation.

## 1. Introduction

Currently human beings are overwhelmed with ever increasing information flows so that one speaks about information overload and information stress (Matitaishvili et al., 2017; Misra et al., 2020). In the era of global digitalization and information expansion, it should be studied how modern neurointerfaces may help to optimize the state of a human being and activate their cognitive resources. Thus, modern cognitive science should develop technologies that could exploit functional brain asymmetry and activity.

### 1.1. Hemispheric asymmetry

Currently the assumption of the dynamic nature of functional hemispheric asymmetry is widely accepted (Floegel & Kell, 2017).

In general, functional hemispheric asymmetry describes functional difference between symmetric brain structures. Once it is developed, it is subject to change as a result of compensatory restructuring of structural functional relations caused by lesions. The asymmetry of functional brain relations can be modulated under different manipulations (Fokin et al., 2004). For example, right-handers may have a greater level of nonspecific activation in the right or in the left hemisphere in a particular state. Some functional states lead to increased elecrophysiological asymmetry up to a statistically significant level. For other functional states, however, such an asymmetry cannot be evidenced. Thus, functional brain asymmetry is determined by a person’s functional state, which is defined as the dynamic nature of functional hemispheric asymmetry (Fokin et al., 2004).

In the current study, we will be referring to the dynamic nature of functional hemispheric asymmetry which is expressed in the asymmetry of hemispheric relations conditioned by the distribution of neuronal activity over symmetrical brain structures in the right and in the left hemisphere.

Other terms that are used to refer to what we call functional hemispheric asymmetry are functional connectivity of hemispheres (Watson et al., 201), asymmetry of hemispheric network topology (Sun et al., 2017), functional hemispheric asymmetries (Floegel & Kell, 2017), hemispheric functional equivalence (Stanković & Nešić, 2020), hemispheric asymmetry of functional brain networks (Cao et al., 2020).

Dynamic brain asymmetry is highly influenced by ‘functional state’ (a term widely used within Russian cognitive science and psychophysiology tradition), which refers to a system response of the body which provides its adequacy to the requirements of the activity (Leonova, 2009). It is shown that the following factors can influence the functional hemispheric asymmetry: mental state (Burdakov, 2010), emotional arousal (Cao et al., 2020), depression (Rotenberg, 2004), anxiety (Adolph & Margraf, 2017), mental fatigue (Sun et al., 2014).

While manual asymmetry characteristics remain more or less constant (Teixeira, 2008; Reiss & Reiss, 2000), functional brain asymmetry characteristics are dynamic across the lifespan. For example, it has been shown that frontal asymmetry is not related to handedness (Schrammen et al., 2020).

One of the ways to quickly and unobtrusively determine functional hemispheric asymmetry is the so-called dichotic listening (Westerhausen, 2019) that generally consists in playing two sound stimuli into two ears. A version of dichotic listening is computer laterometry (Antonets et al., 2011) which is based on dichotic listening and the precedence effect (Tollin, 2003) - the “two-source” lead-lag paradigm (Brown et al., 2015).

This technique consists in the presentation of a series of dichotic stimuli with increasing lead-lag delay duration (Δt). The influence of dichotic stimulation with Δt on the magnitude of the afferent response has been clearly demonstrated in a series of electrophysiological studies of anesthetized cats in which the evoked potentials were recorded from symmetrical points of the posterior hills of the quadriplegium during dichotic stimulation with Δt = 0 and during a 0.1-1.1 ms lead time of one of the ears (Scherbakov & Kosyuga, 1994). Thus, Δt is transformed into the hemispheric asymmetry of excitations. Having reached a certain threshold value, hemispheric asymmetry of afferent excitations acts as an unconditional stimulus for the brain, and it triggers reciprocal relationships between paired nerve centers. As a result of reciprocal relations, efferent flows of one half of the brain are completely suppressed (inhibited), and the other half is weakened (Parenko, 2009). So, although laterometry technology measures subjective sensory space, we can also assess asymmetry of hemispheric relations (distribution of functional activity across hemispheres). The procedure is described in more detail in the Materials and Methods section.

The purpose of this article is to generally explore whether functional hemispheric asymmetry can be changed externally. We will use double biofeedback to induce such a change.

### 1.2. Biofeedback

To start with, the very specifics of neurointerfaces (biofeedback trainings), in which we can holistically (and unobtrusively) influence the organism, the functional state of a person, will be briefly considered.

In general, biofeedback is a method of directional change of the human state in the desired direction. Various studies have shown that biofeedback training can facilitate language acquisition in people with speech disorders (e.g., Adler-Bock et al, 2007; McAllister Byun, 2017; McAllister Byun & Hitchcock, 2012; Preston and Leaman, 2014) as well as in people learning a foreign language (Kartushina et al., 2015).

Biofeedback involves the use of specific hardware-software systems to create realtime visual representations of behavioral or physiological patterns.

Recent research shows that biofeedback is attracting interest not only on the part of physiologists but has also received worldwide approval from specialists in the field of psychology. Such trainings allow to increase the efficiency of human adaptation to various conditions and maximal activation of the inner resources of the human (Balconi et al., 2019). An individual can observe such a mapping and modify his or her behavior to achieve a better fit with the reference model. For example, visual-acoustic biofeedback provides visualization of the spectrum of the acoustic speech signal (e.g., McAllister Byun and Hitchcock, 2012), ultrasonic biofeedback provides real-time visualization of tongue shape and movement within the oral cavity (e.g., McAllister Byun &Hitchcock, 2012), to name but a few.

We suggest that biofeedback can be used as a noninvasive way to change functional hemispheric asymmetry and activity for the following reasons.

1. Biofeedback can reshape neural networks by increasing their connectivity and neuroplasticity (Levine et al., 2000; Villanueva et al., 2011). Globally, biofeedback has been successfully used to alter alpha and gamma brain activity in subjects in order to improve cognitive abilities (Deiber et al, 2020). Biofeedback has demonstrated its effectiveness in ADHD, autism spectrum disorders, substance use, PTSD, and learning difficulties (Niv, 2013). Also, these procedures can result in long-term changes in the distribution of functional activity across the brain (Choi et al., 2011), which allows the application of biofeedback in psychiatric rehabilitation (Markiewcz, 2017).
2. Biofeedback can modify the distribution of functional activity across the cerebral hemispheres.

The technology provides bipolar asymmetry training, during which regulates the distribution of frequencies of different intensity. For example, multidirectional changes in the relative EEG intensity in the brain hemispheres can cause enhancement of cognitive activity and attention processes (Putman, 2005), as well as improvement of executive functions (Kim et al., 2013; Landes et al., 2017). Studies have shown that the effectiveness of biofeedback is determined by many factors such as: motivation (Kleinman, 1981; Saho & Harano, 1984), training (Keefe & Gardner, 1979), respiration (Conde Pastor et al., 2008), and some individual characteristics (Suter, 1979; Li et al., 2019). In general, by unlocking the body’s full potential and mobilizing resources to achieve the goal, one can eliminate excessive psycho-emotional tension and improve the efficiency and rationality of activities not only during biofeedback trainings, but also thereafter.

Most well-studied is biofeedback with the activation of alpha-rhythm. The findings indicate that the ‘alpha state’, along with general relaxation, leads to activation of various types of cognitive activity, in particular – attention (Klimesch, 2012), memory (Nan et al., 2012; van Driel et al., 2012), and even creativity (Fink & Benedek, 2014). After biofeedback training, subjects showed improvements in selective attention (Bauer et al., 2012), error detection (van Driel et al., 2012), and a decrease in anxiety (Bhat, 2010).

### 1.3. Double biofeedback

Double biofeedback is a relatively new technology, which is seen as an effective tool for optimization of the psychophysiological state to realize a wide range of cognitive functions (Fedotchev et al., 2019). The fundamental difference of double biofeedback from the above mentioned one is that there is no need for the subject to intentionally change his (physiological) reactions on his own by some mapping/visualization of them. In this case the loop is closed without conscious participation of the person - but by his/her own physiological reactions. The state is optimized faster, and one session is sufficient for a pronounced effect. The optimal measures for feedback lag are established (Fedotchev et al., 2016). Double biofeedback is technically a more complicated solution, so it is not used as universally as the one-loop biofeedback.

### 1.4. Biofeedback and hemispheric asymmetry

To our knowledge, there are no studies investigating the dynamics of functional hemispheric asymmetry and activity during biofeedback and double biofeedback have been found by the authors of this article. There are some studies investigating the influence of biofeedback on different types of hemispheric asymmetry. The majority of works investigate short-term changes in the asymmetry in alpha-rhythm intensity during biofeedback (Moore, 1984; Zhao et al., 2018; Dziembowska et al., 2016) or frontal asymmetry (John et al., 2001). Separate works are devoted to the study of the asymmetry of slow cortical potentials (Rockstroh et al., 1990). Since functional hemispheric asymmetry can vary depending on the general level of brain activation, which can be caused by prevention of energy depletion and be compensatory during the functional state change (Fokin et al., 2009), such activation during double biofeedback is seen as an efficient tool to ‘manage’ functional hemispheric asymmetry and activity.

The purpose of the current study was to reveal the general patterns of double bio-feedback influence on functional hemispheric asymmetry and activity.

## 2. Materials and Methods

### 2.1. Participants

30 healthy individuals (8 males, 24 females; mean age = 29; SD=8) took part in the current study.

### 2.2. Computer laterometry

To investigate functional brain asymmetry, we used computer laterometry which has been developed based on fundamental neurophysiological research and patented procedures of investigating lateral sensor asymmetry (Scherbakov & Kosyugam 1980; Scherbakov et al., 2003a,b; Scherbakov et al., 2008). Computer laterometry allows to construct various temporal-amplitudinal structures of rectangle sound and noise impulses as well as reaction recording. Stimulus presentation consists in short sound clicks that are given into two ears at registered temporal delays through stereo headphones.

In the current research, the virtual acoustic space consisted in a series of dichotic impulses at a frequency of 3 Hz with the increasing lead-lag delay duration at the rate of 23 microseconds. Sound click intensity was kept constant across participants and did not exceed 40 dB from the monaural hearing threshold with the duration of 50 microseconds.

The procedure was as follows. During the training phase, the participants were familiarized with virtual auditory space stimuli. At the beginning of the experimental phase, the participants were requested to give a joystick response when 1) the sound started shifting from the vertex to one of the ears; 2) the sound reached extreme lateralization, i.e. it was clearly heard around one of the ears; 3) there appeared a wholesome image consisting of two independent sounds in two ears (one dominant and loud sound and the other ‘echo’ sound which is distinct, but quiet). Stimuli were presented first on the left and then on the right-hand side.

To evaluate functional brain asymmetry, basic laterometry parameters were recorded.

1. Δt min L (µs) – lead-lag delay when the virtual auditory space stimulus started shifting from the vertex with left ear advance.
2. Δt min R (µs) – lead-lag delay when the virtual auditory space stimulus started shifting from the vertex with right ear advance.
3. Δt max L (µs) – lead-lag delay at extreme lateralization with left-ear advance.
4. Δt max R (µs) – lead-lag delay at extreme lateralization with right-ear advance.
5. Δt rash L (µs) – lead-lag delay at the ‘echo’-effect with left-ear advance.
6. Δt rash L (µs) – lead-lag delay at the ‘echo’-effect with right-ear advance. Functional hemispheric asymmetry coefficients were calculated as follows.

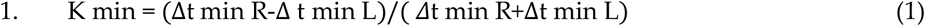

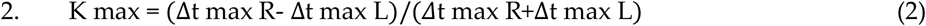

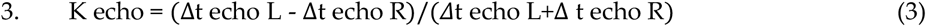

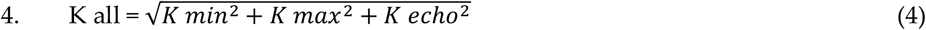

It is claimed that there is a 5 to 1 ratio of contralateral to ipsilateral fibres in the auditory system (Musiek and Baran, 2007). Thus, with the right ear advance we evaluate the functional activity of the left hemisphere with the left ear advance the functional activity of the right hemisphere is estimated (Polevaya, 2009). For K min/max/echo > 0, the right hemisphere is dominant on this parameter. For K min/max/echo < 0, then the left hemisphere is functionally dominant on this parameter.

Based on previous results with regard to tuning functions of multiple neurons at different hierarchical levels of the auditory system and the dynamics of event-related potentials during lead-lag desynchronisation processing, approximate individual modules (Polevaya, 2009) have been defined that are responsible for the three components of the precedence effect (the start time of the virtual auditory space stimulus shifts from the centre, extreme lateralization and ‘echo’, Blauert, 1997; Litovsky et al., 1999; Tollin, Yin, 2003) as well as characteristics of the virtual auditory space stimuli during dichotic simulation with increasing lead-lag delay duration (Δt min, Δt max, Δt echo, Table 1).

**Table 1.**
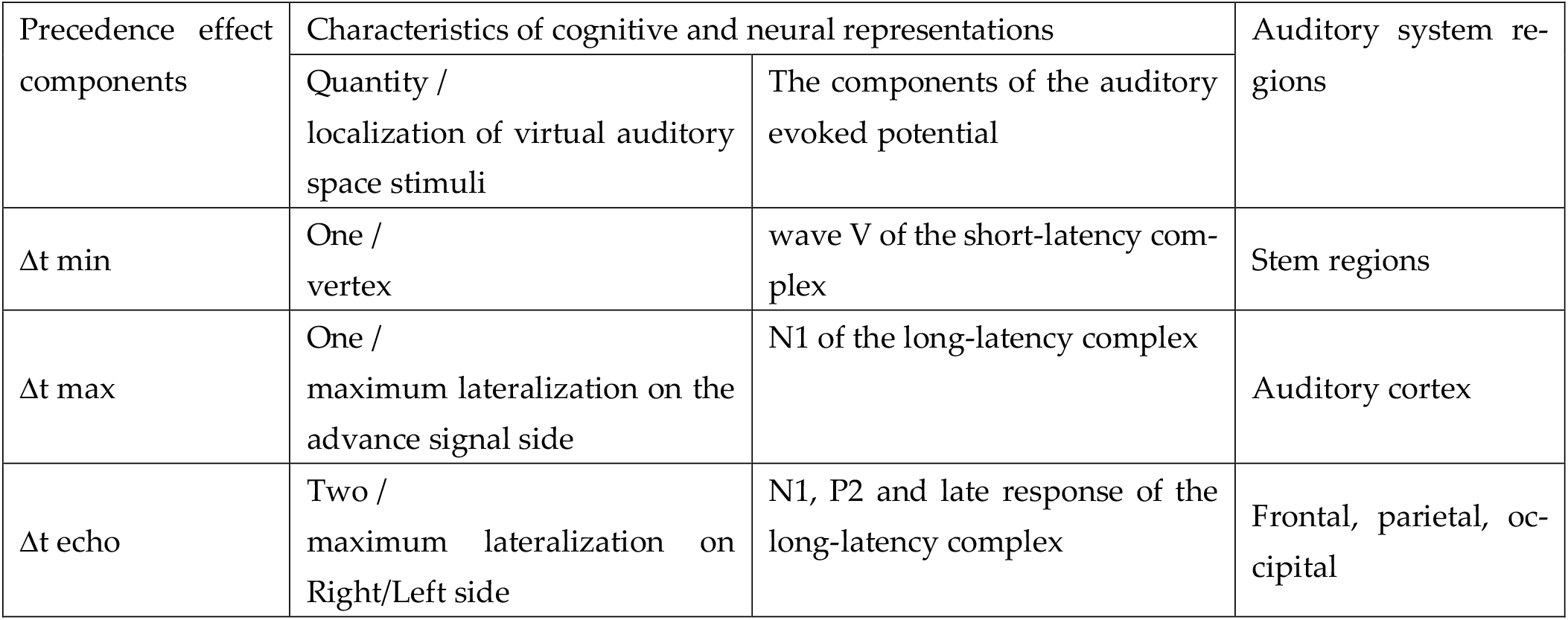
Referencial modules for the precedence effect components

| Precedence effect components | Characteristics of cognitive and neural representations |  | Auditory system regions |
| --- | --- | --- | --- |
|  | Quantity / localization of virtual auditory space stimuli | The components of the auditory evoked potential |  |
| $\Delta t_{\min}$ | One / vertex | wave V of the short-latency complex | Stem regions |
| $\Delta t_{\max}$ | One / maximum lateralization on the advance signal side | N1 of the long-latency complex | Auditory cortex |
| $\Delta t_{\text{echo}}$ | Two / maximum lateralization on Right/Left side | N1, P2 and late response of the long-latency complex | Frontal, parietal, occipital |

For the first precedence effect component, the lead-lag delay is measured that is sufficient for the perception of the fusion sound to have shifted from the vertex (the centre of the inter-ear bend) to the side of advance stimulus presentation (Δt min). That is, lead-lag duration reflects how far the advance hemisphere is labile, i.e., it can be pre-activated prior to dominating.

For the first precedence effect component, the lead-lag delay that provides maximal lateralization of the virtual auditory space stimulus (Δt max) is measured, that is, the shift towards the extreme side location equivalent to the monaural localisation. Thus, Δt max reflects to what extent the advance hemisphere is excitable, i.e., how quickly it can start dominating.

For ‘echo’, lead-lag delay is measured that is sufficient to transform two temporally disparate signals into two spatially disparate virtual auditory space stimuli. That is, Δtrash reflects to what extent the advance hemisphere is stable, i.e., how long it can preserve dominance and inhibit the other hemisphere.

Thus, basic laterometry parameters (Δt min, Δt max, Δt echo) are related to hemisphere lability, excitability, and stability.

The lower the Δt min the higher the lability of the hemisphere that is opposite to the sound shift direction, which reflects lower activation threshold for neuronal corollaries in the brain stem.

The lower the Δt max, the greater the excitability of the hemisphere that is opposite to the sound shift direction, which reflects lower activation threshold for neural corollaries in the primary auditory cortex.

The lower the Δt echo, the lower the stability of the hemisphere that is opposite to the sound shift direction, which reflects shorter time span of neuronal activity in the frontal, parietal, and occipital cortex.

The values Δt min, Δt max, Δt echo for the left and right sound shift (to determine the dominance of the right and the left hemisphere) obtained as a result of laterometry allow to evaluate functional brain asymmetry in terms of lability, excitability and stability.

Thus, the following two phenomena were looked at with the help of computer laterometry in the current study.

1. Functional hemispheric asymmetry, as reflected in K min, K max, K echo.
2. Functional hemispheric activity, as reflected in Δt min, Δt max, Δt echo and interpreted as lability, excitability, and stability.

### 2.3. Double biofeedback

Double biofeedback is a dual loop biofeedback system for monitoring rhythmic brain activity (with vertex electrode) and correcting/optimizing cognitive functions and emotional state. Unlike similar techniques, this technology allows detection of spectral components of brain biopotentials with high frequency resolution, and in combination with the original resonance stimulation methods gives an opportunity to detect and analyze individual narrow-frequency EEG oscillators of an individual (Fedotchev et al., 2016; Fedotchev et al., 2019).

This device eliminates the limitations of existing EEG-based biocontrol methods through unique innovations. First, it does not use predetermined, overly broad-band traditional EEG rhythms (theta - 4-8 Hz, alpha - 8-13 Hz, beta - 13-25 Hz, etc.), but automatically detects in real time, narrow-frequency EEG oscillators that are characteristic of a particular individual and significant for them. The promising nature of this approach is related to the functional heterogeneity of EEG and the effectiveness of using narrow EEG frequencies of the subject in EEG biofeedback (Hammond, 2011). Secondly, it facilitates teaching a person to self-regulate his or her state by introducing an additional feedback loop that works automatically simultaneously with the conscious adaptive biocontrol loop. Introduction of automatic modulation of sensory influences by endogenous rhythms eliminates dependence of the procedure’s efficiency on the subject’s motivation level. Thirdly, inclusion of two feedback channels and an EEG channel into the feedback loop will provide faster influence on the functional activity of hemispheres. The mentioned advantages of the technology, providing its increased efficiency, are implemented in a microprocessor device, which was applied in this study.

### 2.4. Study design

The experiment was administered in several consequent steps.

1. The participant was asked to wear headphones and LED glasses. The headphones were used to laterometry as well as to play double biofeedback sound stimuli.

2. The evaluation of functional hemispheric asymmetry and activity with the help of laterometry.

3. Double biofeedback

3.1 The recording of narrow-frequency components withing the pre-defined EEG range (4-20 Hz with 0.1 Hz frequency increase every three seconds) that are dominant for a particular participant. Total recording duration was equal to 480 seconds.

3.2 General double biofeedback mode where the current amplitude of the respective EEG oscillator from the pre-defined range (4-20 Hz) is transformed into feedback sound signals and the current amplitude of the respective alpha EEG oscillator is used to modulate the intensity of sinusoidal light signals that are generated at the frequency rate of this oscillator. Total recoding duration was equal to 300 sec.

4. The evaluation of the functional hemispheric asymmetry and activity with the help of laterometry.

The study design and procedure were approved by the Ethics Committee of Lobachevsky State University, and all participants provided written informed consent in accordance with the Declaration of Helsinki.

### 2.5. Data analysis

To estimate the difference between the parameters of functional brain activity (Δt min, Δt max, Δt echo) before and after double biofeedback, as well as during the sound shift to the left and to the right, T-test for dependent samples was used. To analyze proportional differences two proportion Z-Test was used.

To estimate the differences on the investigated parameters for male and female participants, non-parametric Mann-Witney test for independent samples was used. Pivot tables were generated in MS Excel v 2102.

## 3. Results and discussion

### 3.1. Functional hemispheric asymmetry dynamics

The analysis of functional hemispheric activity was performed along three lines: 1) hemispheric dominance (by lability, excitability, stability) before and after the biofeedback; 2) statistical significance of functional brain asymmetry coefficient change by lability, excitability and stability; 3) personal analysis of the dynamics of hemispheric activity dominance as to lability, excitability, stability.

The distribution of hemispheric dominance by lability, excitability, and stability before and after the biofeedback is shown in Figure 1 (a, b, c).

**Figure 1.**
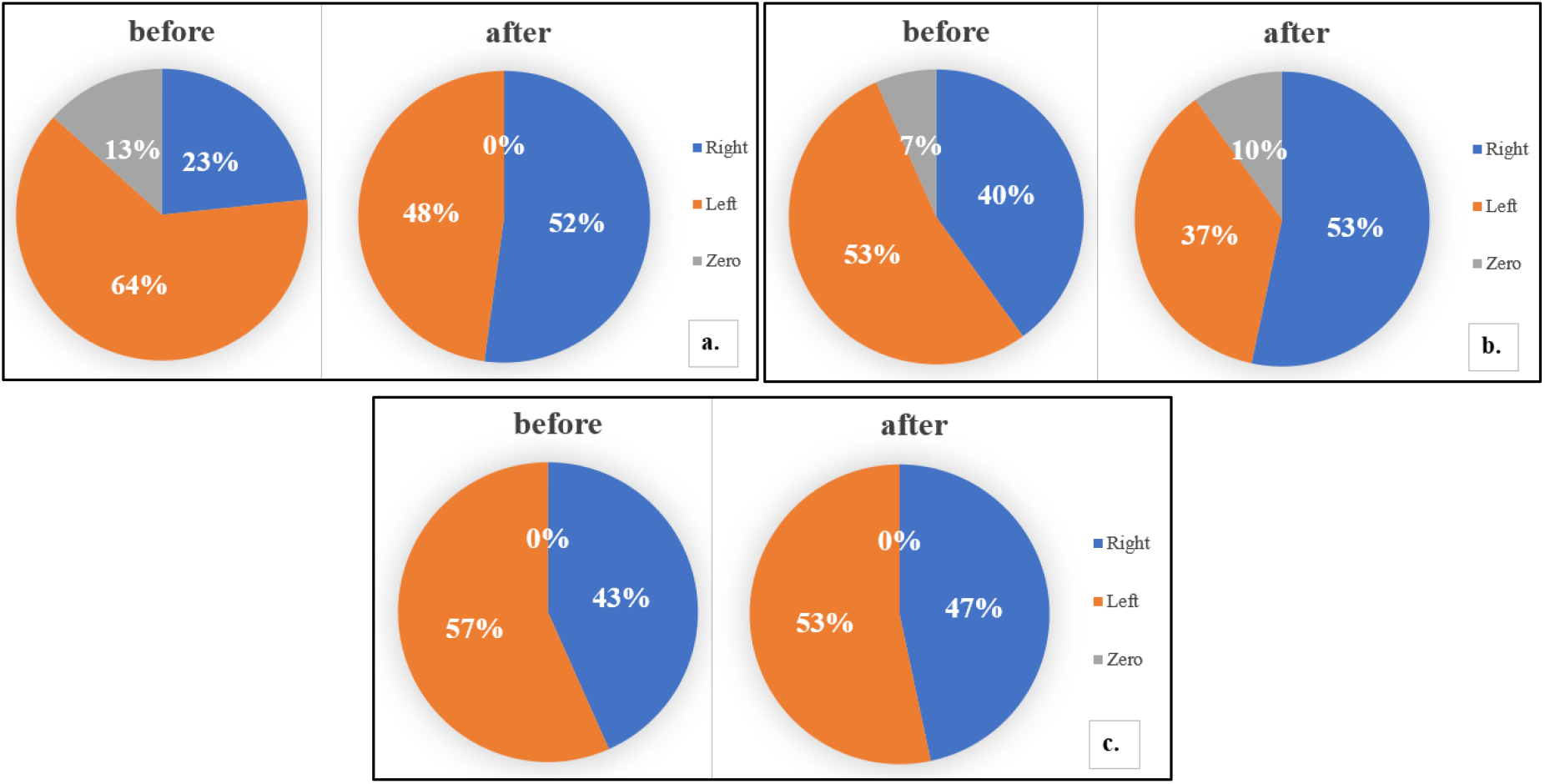
Dominant hemisphere by lability (a), excitability (b), and stability (c) before and after biofeedback (‘Right’ - K min/max/echo > 0, ‘Left’ - K min/max/echo < 0, ‘Zero’ - K min/max/echo = 0)

The sample is relatively evenly distributed by functional activity dominance in terms of excitability and stability both before and after double biofeedback. As to lability, the majority of participants were left hemisphere dominant prior to double feedback (19 participants – 64 %). Their prevalence is statistically significant (Two Proportion Z-Test (p<0.01)). Such an effect is still to be explained. It can be connected with predominantly evening testing sessions, yet the influence of the time of testing has not been previously investigated and remains to be investigated in the future.

The results of statistical analysis of functional brain asymmetry coefficients before and after feedback are presented in Table 2.

**Table 2.** The functional hemispheric asymmetry coefficients before and after double biofeedback training (means and standard deviations), t-test and p-values.

|  | Mean |  | Std.Dv. |  | t | p |
| --- | --- | --- | --- | --- | --- | --- |
|  | before | after | before | after |  |  |
| K min (lability) | -0.064 | -0.018 | 0.136 | 0.136 | -1.3 | 0.197 |
| K max (excitability) | -0.011 | 0.011 | 0.098 | 0.099 | -0.9 | 0.372 |
| K echo (stability) | -0.020 | 0.005 | 0.125 | 0.096 | -0.9 | 0.373 |
| K all | 0.203 | 0.179 | 0.078 | 0.070 | 1.1 | 0.281 |

Functional hemispheric asymmetry coefficients did not change before and after the double biofeedback training. So, globally functional brain asymmetry did not change as to any of the components (lability, excitability or stability). It can be rooted in more complex mechanisms underpinning brain activity during double biofeedback training. Secondly, it can be caused by other factors that have not been accounted for in the current study.

We then conducted personalised analysis of functional hemispheric asymmetry change for the whole sample. The data are summarized in Table 3.

**Table 3.**
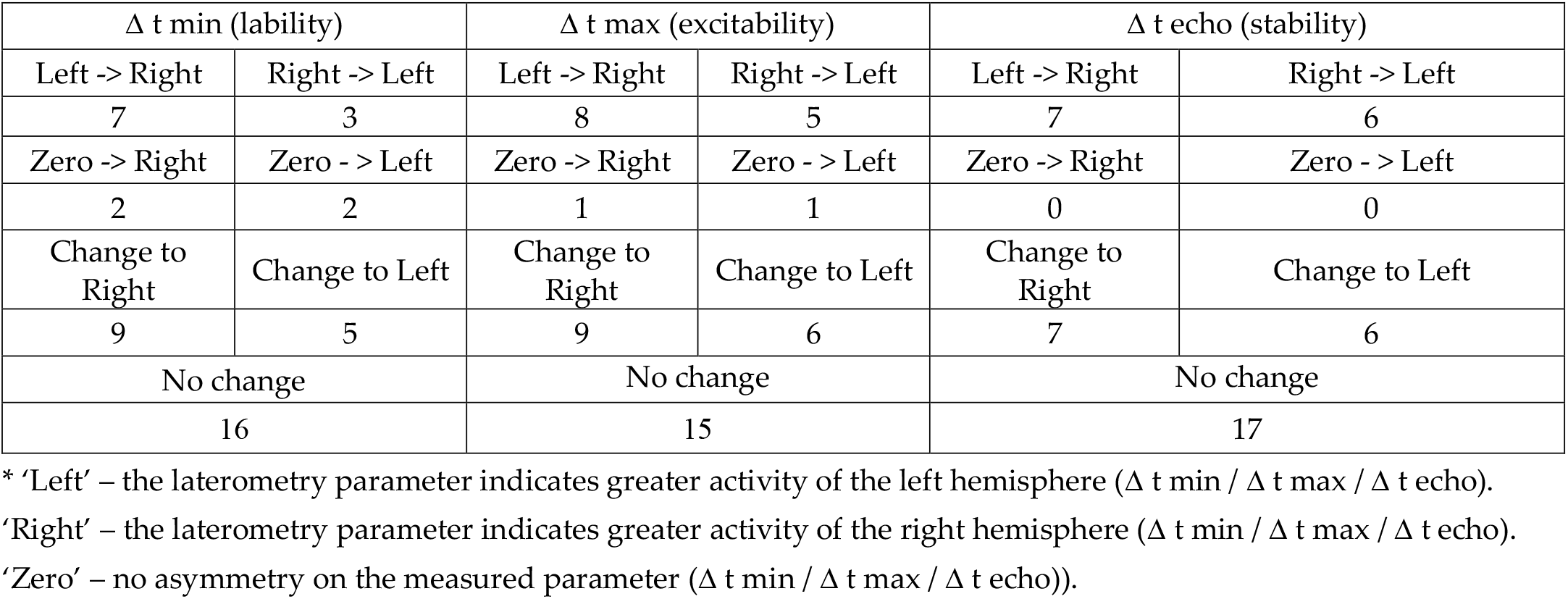
Changes in the dominance of hemispheric activity by lability, excitability, stability*

Half of the participant sample did not experience any functional hemispheric activity change on any of the components – lability, excitability, stability (53%, 50%, 57% respectively). Among those who have experienced asymmetry change, there is a tendency to the rightward shift. This tendency is more pronounced for lability. Among 14 people who experienced asymmetry inversion, in 9 (64%) it changed to the right, yet this change is not statistically significant (Two Proportion Z-Test p = 0.14). Thus, a greater sample is needed to prove this tendency.

In general, functional hemispheric asymmetry change in the current experiment supports previous claims with regard to its dynamic nature and its possible modification under the influence of different factors.

### 3.2. Functional hemispheric activity dynamics

The analysis of functional hemispheric activity was performed along two lines: 1) the direction of the sound shift (Left vs Right); 2) measurement stage (before vs after double biofeedback)

No difference in Δ t max and Δ t echo was observed for sound shift to the left or to the right at all stages of the experiment. Only one statistically significant difference was observed in Δ t min before double biofeedback – see fig. 2.

**Figure 2.**
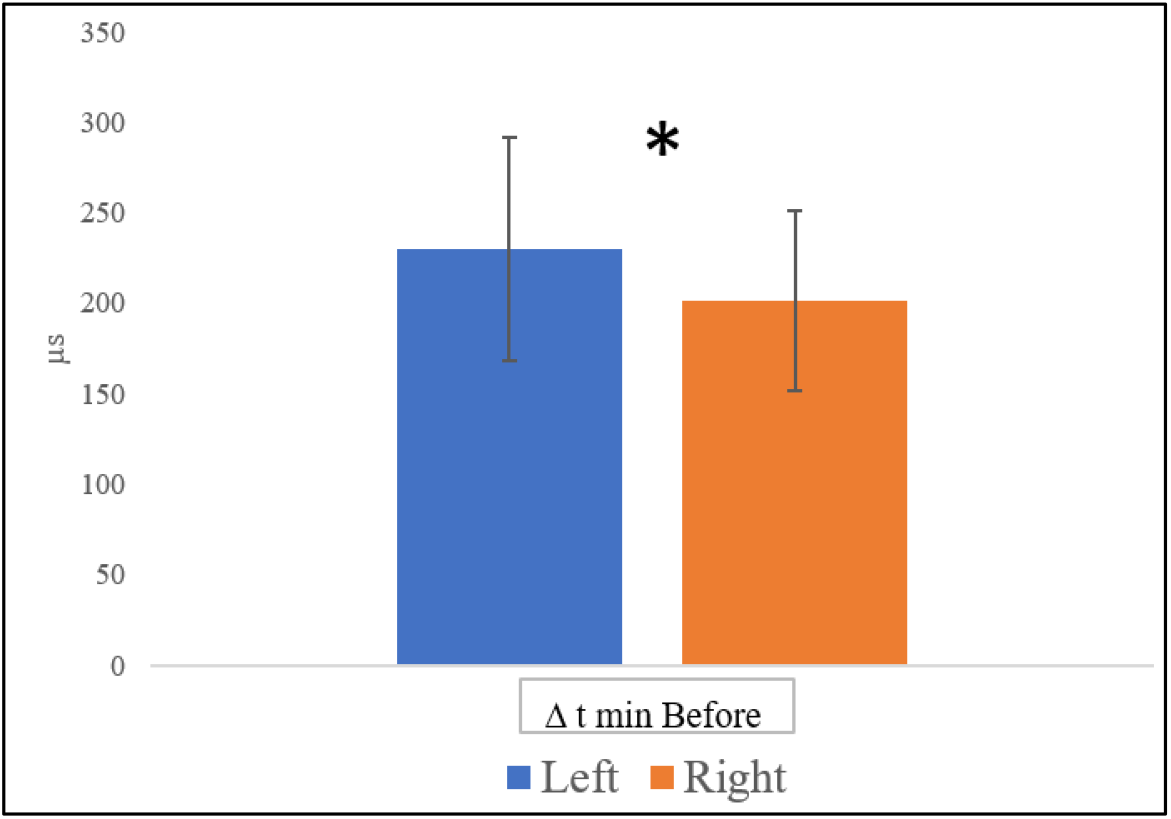
Mean values for Δ t min Left and Right before double biofeedback (vertical bars denote standard deviations; * - p<0.05 – T-test for dependent samples)

Thus, prior to double feedback there was a statistically significant difference in hemispheric lability: Δ t min before Left is greater than Δ t min before Right (t=2.6, p<0.05). It is in line with the general distribution of the sample on initial functional hemispheric asymmetry – see fig. 1a.

Δ t min Left, Δ t echo Left, and Δ t echo Right were significantly reduced after double feedback training - see fig. 3.

**Figure 3.**
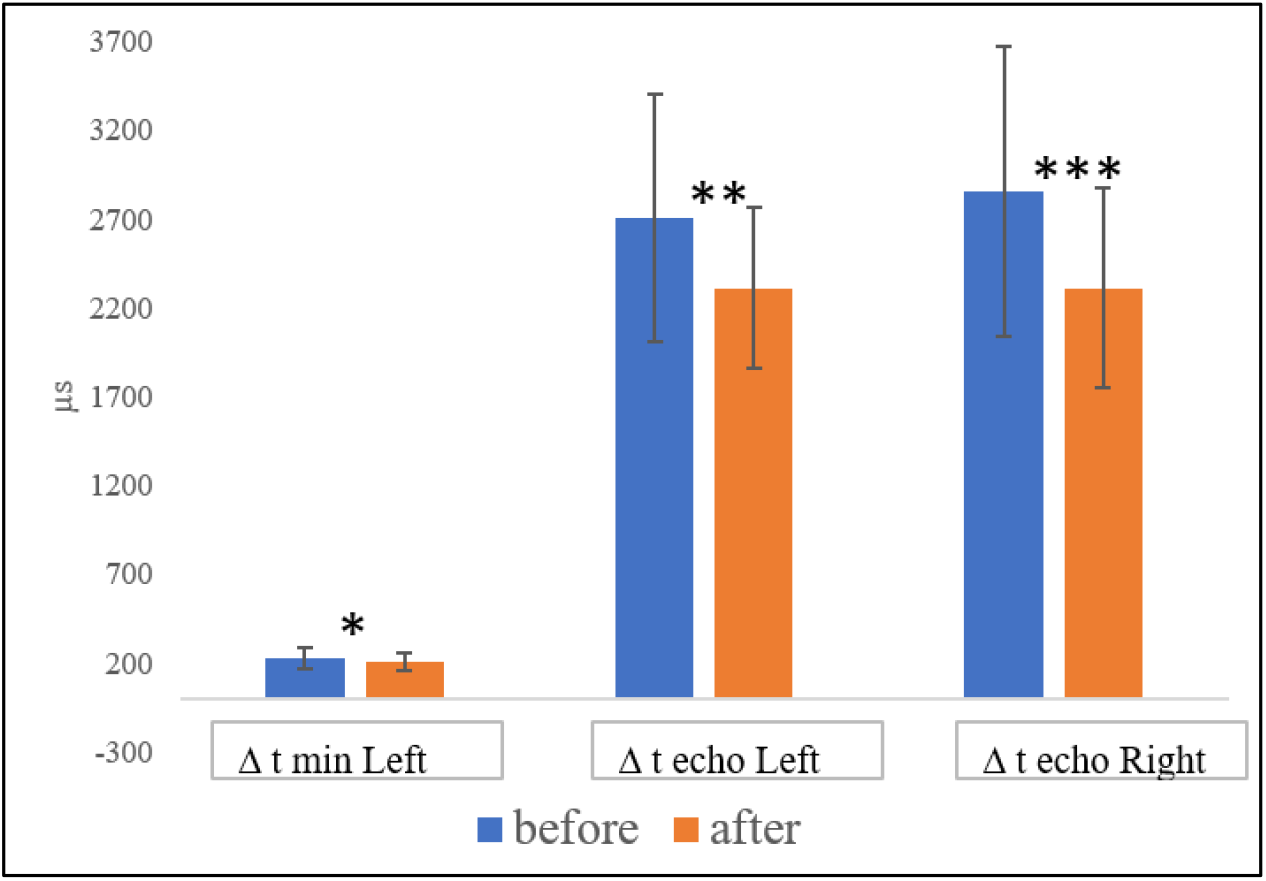
Mean values for Δ t min Left, Δ t echo Left, and Δ t echo Right before and after double biofeedback (vertical bars denote standard deviations; * - p<0.05, * * - p<0.01, * * * - p<0.001 – T-test for dependent samples)

Thus, we found that there was a statistically significant acceleration of information processing in the brain stem of the right hemisphere as well as in the frontal, parietal and occipital lobes of the right and the left hemispheres.

Stability reduction reveals that one of the hemispheres is quick to give preferential processing of sound information over to the other hemisphere. Possibly, double biofeedback contributes to the deactivation of reciprocal inhibition. One can say that double bio-feedback contributes to general brain activation in quite a number of dimensions, which is in line with what has been shown previously (Klimesch, 2012; Nan et al., 2012; van Driel, 2012; Fink & Benedek, 2014; Bauer 2012; Bhat, 2010).

Such an effect may seem to be related to timing or learning. Yet, functional brain asymmetry evaluation with the help of computer laterometry was also used in other projects in a variety of contexts. The tendency to lability and stability reduction has not been reported (e.g., Demareva &Polevaya, 2014). Future research should take it into account by introducing a control group of participants whose functional hemispheric asymmetry is screened at the beginning and at the end of the double biofeedback time window. Another non-biofeedback treated control group who is just listening to music and experiencing LED flashes which are not modulated by EEG is also needed.

It is also worth mentioning that Mann-Witney test did not show any significant differences between male and female participants on any of the parameters. To study this aspect further, a gender-balanced sample is needed. The goal of the current pilot investigation was to trace basic tendencies in hemispheric asymmetry and activity dynamics before and after double biofeedback training. Gender difference in functional brain asymmetry in its various manifestations is being widely discussed in the literature (see Hirnstein et al., 2019 for review).

## 4. Conclusions

1. Double biofeedback did not alter functional brain asymmetry on any of its characteristics – lability, excitability, or stability.
2. An increase in the right hemisphere lability and a decrease in the stability of both hemispheres was evidenced after double biofeedback training.
3. Further research on double biofeedback is needed with different control groups, varying experimental conditions in treatment groups as well as controlling for gender, age, and time of testing.

## Author Contributions

“Conceptualization and methodology Valeriia Demareva; investigation, Elena Mukhina and Valeriia Demareva; analysis Valeriia Demareva, writing—original draft preparation Valeriia Demareva and Elena Mukhina; writing—review and editing Tatiana Bobro.

## Funding

This research was funded by Lobachevsky State University, grant number H-456-99_20-2 (‘Evaluation of the effectiveness of neurobiofeedback to optimize the functional state in learning a foreign language’).

## Acknowledgements

The authors would like to thank Alexander Fedotchev and Alexander Bondar for providing the double biofeedback device and Sofia Polevaya for explaining the specifics of the device in the context of this study.

## Conflicts of Interest

The authors declare no conflict of interest.

## Notes

### Competing Interest Statement

The authors have declared no competing interest.

